# Up and to the light: intra- and interspecific variability of photo- and geo-tactic oviposition preferences in genus *Trichogramma*

**DOI:** 10.1101/2021.03.30.437671

**Authors:** V. Burte, G. Perez, F. Ayed, G. Groussier, L. Mailleret, L. van Oudenhove, V. Calcagno

## Abstract

*Trichogramma* are parasitic microwasps much used as biological control agents. The genus is known to harbor tremendous diversity, at both inter- and intra-specific levels. The successful selection of *Trichogramma* strains for biocontrol depends on characterizing the existing diversity in this group, especially regarding oviposition performance and behavior. Oviposition preferences in relation to environmental cues such as light and gravity have received little attention so far, although they are suspected to play an important role in microhabitat selection and parasitism patterns. The extent of their variability, and their potential correlated variations, is virtually unknown. Here we use a novel experimental approach relying on automatic image analysis to characterize the oviposition preferences in relation to light and gravity, as well as their interaction, in 25 populations of *Trichogramma* from five species. We show that most *Trichogramma* populations and species harbour preferences for light and preferences for elevated parts. However, the two traits harbor significant inter and intraspecific variation. The effects of light and gravity on oviposition patterns were found to be almost perfectly additive overall, with two exceptions. Oviposition preference patterns were not static but very plastic in time: preferences tended to relax over consecutive days, and the strongest preferences relaxed the fastest, presumably because of the density-dependent effect of resource depletion. A correlation of oviposition patterns with the vegetation stratum at which populations were sampled suggests that different species/populations may be associated with different strata with corresponding differentiation in light- and gravity-related oviposition preferences.

This article has been peer-reviewed and recommended by *Peer Community* in Zoology doi: https://doi.org/10.24072/pci.zool.100008

## Introduction

*Trichogramma* wasps are among the insects most used as biocontrol agents for crop protection. They are able to control more than 30 pests, mostly from Lepidoptera (Consoli et al. 2010). In France for instance, more than 120,000 hectares of maize fields (about 25% of the overall surface protected against the European Corn Borer) are treated using mass releases of industrially produced *Trichogramma brassicae. Trichogramma* wasps are tiny parasitic wasps that lay their eggs into the eggs of their insect hosts. Several aspects of their ecology, physiology and diversity are still relatively little understood, and estimates of natural population densities or genetic structure are virtually lacking, owing to the difficulty of sampling and studying natural populations. Only a few species and strains have been studied with attention and even fewer are commercially produced and used for crop protection. Considering that more than 200 species are documented in the genus (Consoli et al. 2010), probably with ample geographic variation and/or cryptic species, there is great scope for selecting and using new strains of *Trichogramma* in biocontrol programs. This, however, requires an important phenotyping effort to characterize the variability across populations and species. The task is made complicated, in addition to the imperfect taxonomic knowledge of the group, by the minute size of these insects (less than a half millimeter).

It is generally recognized that the efficiency of *Trichogramma* as biocontrol agents is determined by classical life-history traits such as cold tolerance, lifespan or fecundity (van Lenteren et al. 2003). Beyond these, more complex traits such as movement strategies and dispersal are increasingly recognized as important predictors of field performance and optimal deployment, and have consequently received significant attention in the literature (e.g. Bigler et al. 1988; van Lenteren et al. 2003; Kölliker-Ott et al. 2004; Fournier et al. 2005; Lartigue et al. 2020). But further dimensions of *Trichogramma* phenotypic traits, in particular with respect to micro-habitat selection and preference for ecological variables, remain largely understudied. This is unfortunate, for it is important to understand how *Trichogramma* species explore their environment and distribute themselves in relation to ecological variables. This is in particular important if we are to predict how different strains or combination of species would perform in particular settings and to make sense of the results of biocontrol assays. A primary determinant of the success of biocontrol programs is indeed the selection of parasitoid strains that fit the environment conditions they will be released in (see Gariepy et al 2015).

One of the simplest possible example is microhabitat selection with respect to light and to gravity. It is generally thought that *Trichogramma* wasps, like many similar insect species, possess a natural preference for light and an innate tendency to climb upward. However, the effect of these orientation cues are rarely rigorously studied, nor is their actual impact on patterns of host parasitism on the longer term. Similarly, their variation across species and populations, and their association with habitat characteristics or potential implication in ecological differentiation remain to be investigated in *Trichogramma*. Such traits have been found to be important for adaptation and host specialization in a number of insect species (e.g. Brodeur & McNeil 1990; Hunt & Nault 1991; Calcagno et al. 2010; Singer 2015).

Within *Trichogramma*, there are clear indications that oviposition preferences with respect to light and gravity exist and are variable across species. The fact that *Trichogramma* species exhibit positive phototaxis and/or negative geotaxis is often taken for granted, and sometimes used for practical purposes, when developing experimental setups or rearing protocols (e.g. Honda et a. 1999; Ambrosius et al. 2005; Henderson et al. 2015). Regarding attraction to light, *T. evanescens* females are indeed found to be attracted to light-traps (Brower & Cline 1984), and, more recently, van Atta et al. (2015) observed that *T. spp* preferred to oviposit in areas more exposed to UV-B light, with consequences on resulting parasitic success. With respect to geotactic preferences, Thorpe (1985), reported that species *T. pretiosum* and *T. minutum* differed in the height at which they preferentially parasitized eggs, even though the underlying behavioral mechanism remained unknown. This difference was hypothesized to explain the relative parasitism performance of the two species in different settings (Thorpe & Dively 1985). Similar patterns of vertical oviposition preference (e.g. Smith 1988; Grieshop et al. 2014) or preferences between upper/lower parts of leaves (Gardner & Hoffmann 2020) are often observed in *Trichogramma*, but, again, the actual contributions of photo- or geo-orientation are usually left hypothetical. Explicit studies of these traits remain extremely scarce.

In this article, we investigate the patterns of photo- and geo-tactic oviposition preference in *Trichogramma*. Taking advantage of an important sampling of natural *Trichogramma* populations in Western Europe (Ris et al. 2018), and on novel phenotyping methods, we were able to compare 25 populations from five species, and to study the two traits individually as well as in interaction, over several days of parasitism. We could thus test whether (i) *Trichogramma* species showed systematic differences in their photo/geo-tactic preferences; (ii) there existed variation in these traits among different populations from the same species; (iii) photo- and geo-tactic preferences were distributed independently or showed consistent associations. In addition, using data on the vegetation stratum at sampling, we could evaluate whether populations from different strata showed consistent differences in their oviposition preference traits.

## Material and Methods

### *Trichogramma* species and populations

We compared 25 populations of *Trichogramma* belonging to five classical Paleoartic species: six from *T. cacoeciae*, five from *T. cordubensis*, four from *T. evanescens*, four from *T. principium*, and six from *T. brassicae. Trichogramma cacoeciae* and *T. cordubensis* are thelytokous, while the others are arrhenotokous. Species identities were assessed using morphological and molecular markers, and within the six *brassicae* populations, four were identified as belonging to a specific genetic cluster called *brassicae – euproctidis*, while the other two belonged to the main *brassicae* cluster. Populations were sampled in France or Spain, using sentinel *Ephestia kuhniella* eggs on different host plants, and established in the lab a few years prior to experiments, or occasionally longer. All 25 strains are maintained in, and available from the EP-Coll Biological Resource Center (Ris et al. 2018) in Sophia-Antipolis upon request. Further information on these strains can be found in Table 1.

**Table 1:**
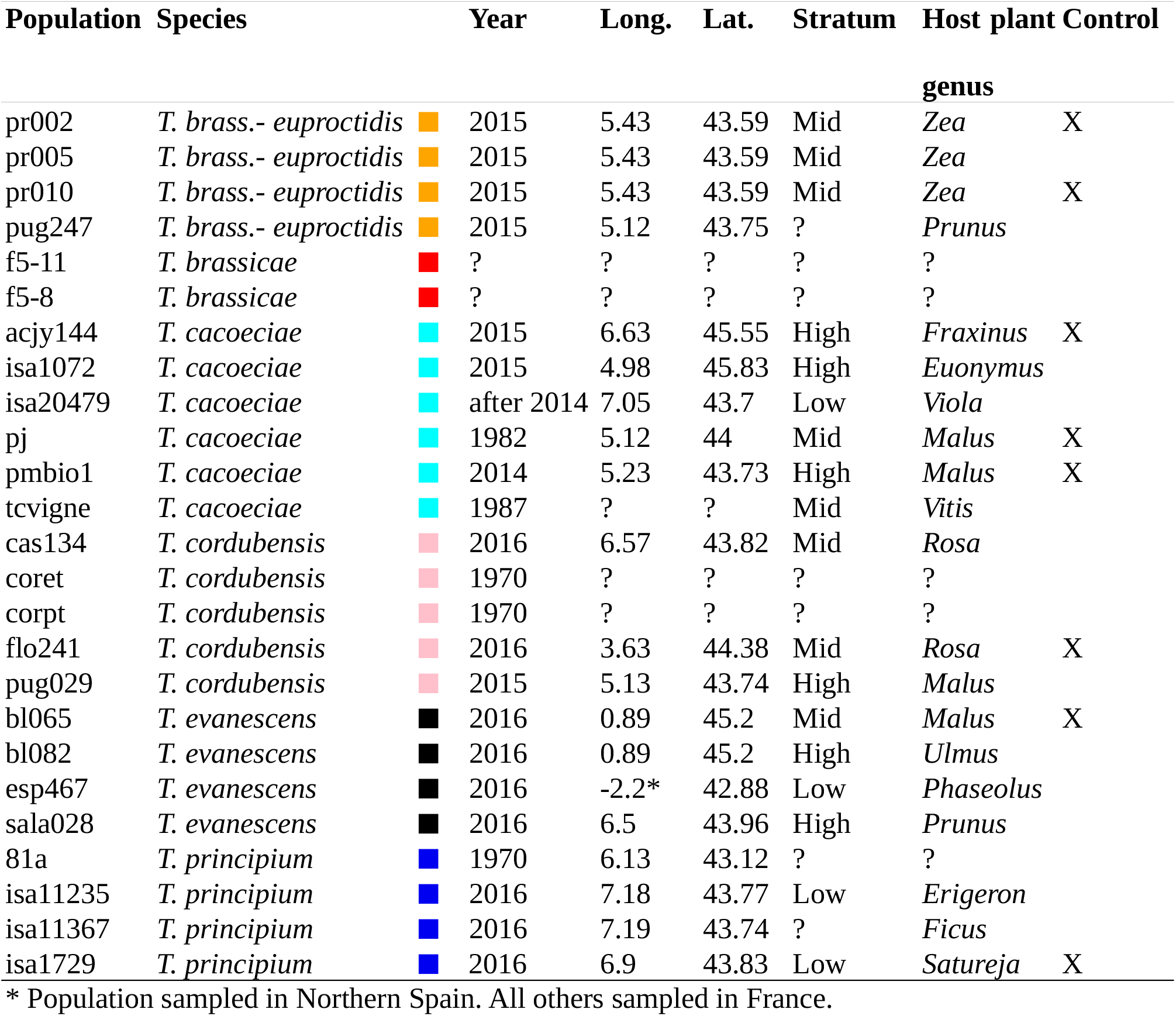
List of the 25 populations studied. For each population, species status, year and location of sampling are provided when available. Stratum refers to the height class at which the sampling was realized, and host plant genus refers to the host plant on which the sentinel eggs were attached. Control indicates whether a control treatment was performed for this population (see Methods).

| Population | Species |  | Year | Long. | Lat. | Stratum | Host plant<br>genus | Control |
| --- | --- | --- | --- | --- | --- | --- | --- | --- |
| pr002 | *T. brass.- euproctidis* | ■ | 2015 | 5.43 | 43.59 | Mid | *Zea* | X |
| pr005 | *T. brass.- euproctidis* | ■ | 2015 | 5.43 | 43.59 | Mid | *Zea* |  |
| pr010 | *T. brass.- euproctidis* | ■ | 2015 | 5.43 | 43.59 | Mid | *Zea* | X |
| pug247 | *T. brass.- euproctidis* | ■ | 2015 | 5.12 | 43.75 | ? | *Prunus* |  |
| f5-11 | *T. brassicae* | ■ | ? | ? | ? | ? | ? |  |
| f5-8 | *T. brassicae* | ■ | ? | ? | ? | ? | ? |  |
| acjy144 | *T. cacoeciae* | ■ | 2015 | 6.63 | 45.55 | High | *Fraxinus* | X |
| isa1072 | *T. cacoeciae* | ■ | 2015 | 4.98 | 45.83 | High | *Euonymus* |  |
| isa20479 | *T. cacoeciae* | ■ | after 2014 | 7.05 | 43.7 | Low | *Viola* |  |
| pj | *T. cacoeciae* | ■ | 1982 | 5.12 | 44 | Mid | *Malus* | X |
| pmbio1 | *T. cacoeciae* | ■ | 2014 | 5.23 | 43.73 | High | *Malus* | X |
| tcvigne | *T. cacoeciae* | ■ | 1987 | ? | ? | Mid | *Vitis* |  |
| cas134 | *T. cordubensis* | ■ | 2016 | 6.57 | 43.82 | Mid | *Rosa* |  |
| coret | *T. cordubensis* | ■ | 1970 | ? | ? | ? | ? |  |
| corpt | *T. cordubensis* | ■ | 1970 | ? | ? | ? | ? |  |
| flo241 | *T. cordubensis* | ■ | 2016 | 3.63 | 44.38 | Mid | *Rosa* | X |
| pug029 | *T. cordubensis* | ■ | 2015 | 5.13 | 43.74 | High | *Malus* |  |
| bl065 | *T. evanescens* | ■ | 2016 | 0.89 | 45.2 | Mid | *Malus* | X |
| bl082 | *T. evanescens* | ■ | 2016 | 0.89 | 45.2 | High | *Ulmus* |  |
| esp467 | *T. evanescens* | ■ | 2016 | -2.2* | 42.88 | Low | *Phaseolus* |  |
| sala028 | *T. evanescens* | ■ | 2016 | 6.5 | 43.96 | High | *Prunus* |  |
| 81a | *T. principium* | ■ | 1970 | 6.13 | 43.12 | ? | ? |  |
| isa11235 | *T. principium* | ■ | 2016 | 7.18 | 43.77 | Low | *Erigeron* |  |
| isa11367 | *T. principium* | ■ | 2016 | 7.19 | 43.74 | ? | *Ficus* |  |
| isa1729 | *T. principium* | ■ | 2016 | 6.9 | 43.83 | Low | *Satureja* | X |

\* Population sampled in Northern Spain. All others sampled in France.

The choice of these populations was dictated by availability, but sought to cover the available variability (both in terms of species and geographic origins within species). We restricted ourselves to Palearctic species that had been sampled using the same protocol at at about the same time, so as to make them as comparable as possible. We also focused on species that are found to be most abundant in the field and/or have interest for biocontrol programs. Most populations had been sampled one to three years before the experiments (ca. 12-35 generations), but we also included some populations that had been maintained in the lab for much longer (up to 25 years; see Table 1), to assess if they differed in any systematic way. Indeed, evolution in the laboratory might affect genetic composition and phenotypic traits. Rearing conditions, in which host eggs (UV-sterilized *E. kuhniella* eggs obtained from the Bioline company) are plentiful and distributed uniformly, within tubes maintained horizontally in a well-lit incubator, are not expected to select for a particular taxis. They could on the contrary favor the loss of existing taxies. For 18 of the 25 populations, we also know the vegetation stratum from which they were sampled, depending on the height at which the sentinel eggs were located and the type of host plant, as categorized in three classes: “Low” (<30 cm above ground), “Mid” (30cm-1m) or “High” (>1m; see Table 1).

Prior to experiments, populations were placed in the same incubator for 2-3 generations, with a 12:12 LD photoperiod and a constant temperature of 25°C, switching to 19°C after a few days, in order to set generation times at exactly 14 days. This allowed to synchronize different populations and facilitated experimentation.

### Experimental protocol

Individuals in each population were synchronized so that their emergence could be predicted to occur on Wednesdays, with a precision of about 24 hours. The day before that predicted emergence, parasitized *E. kuhniella* eggs, recognizable from their dark colour, were taken from the rearing tubes and placed into experimental tubes. Those were cylindrical 15cm-long glass tubes (1cm of inner diameter) containing a 13cm long piece of thin cardboard, and sealed on both sides with an organic cotton plug. The piece of cardboard was covered with 10 circular patches of fresh *E. kuhniella* eggs, each containing about 80 eggs, and evenly distributed along the cardboard, with a 1cm spacing between them (see Figure 1a). Eight parasitized eggs (forming the *Trichogramma* inoculum) were placed in the middle of the Individuals in each population were synchronized so that their emergence could be predicted to occur on Wednesdays, with a precision of about 24 hours. The day before that predicted emergence, parasitized *E. kuhniella* eggs, recognizable from their dark colour, were taken from the rearing tubes and placed into experimental tubes. Those were cylindrical 15cm-long glass tubes (1cm of inner diameter) containing a 13cm long piece of thin cardboard, and sealed on both sides with an organic cotton plug. The piece of cardboard was covered with 10 circular patches of fresh *E. kuhniella* eggs, each containing about 80 eggs, and evenly distributed along the cardboard, with a 1cm spacing between them (see Figure 1a). Eight parasitized eggs (forming the *Trichogramma* inoculum) were placed in the middle of the cardboard piece, thus sitting exactly in between the two innermost egg patches. Two drops of water-diluted honey were also placed next to them. All eggs were glued on the cardboard with water-diluted solvant-free glue. The cardboard used was pink in color to maximize contrast with host eggs and facilitate image analyses (next Section). The tubes were then disposed in a closed room at a constant 24°C and 75% humidity. Tubes were let there for the duration of an experiment, i.e. five days, from Wednesday to Sunday. During that period, adult *Trichogramma* emerged from the parasitized eggs introduced in the middle of the tubes, mated (for sexual species), searched for fresh eggs and parasitized them, and eventually died. The exact numbers of *Trichogramma* females emerging thus varied among tubes, theoretically lying between 0 and slightly more than eight, causing variations in the total rate of parasitism.

**Figure 1:**
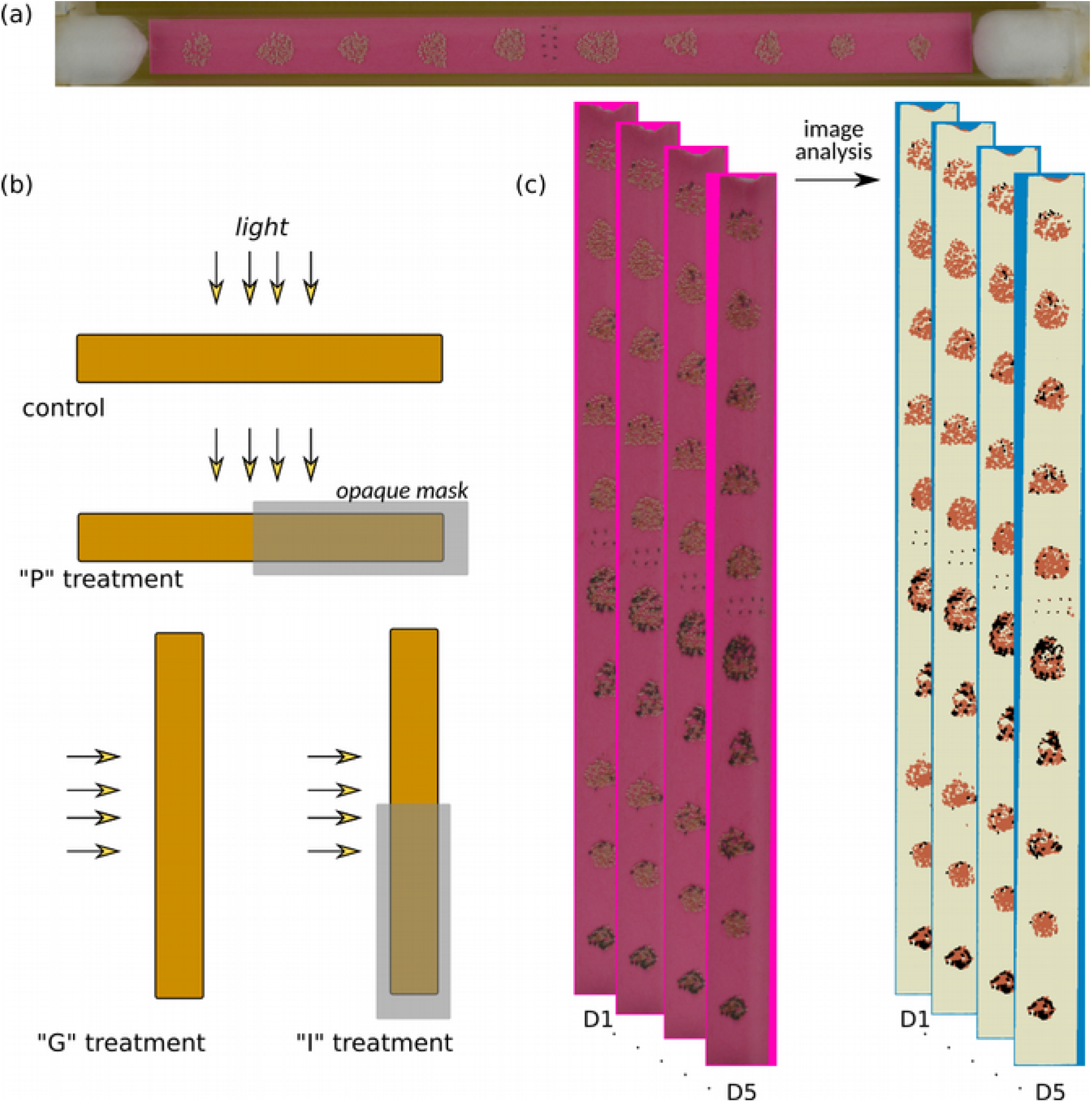
Experimental protocol and image analysis technique. (a) An experimental tube, containing 10 patches of host eggs, evenly spaced along its length. Eight ready-to-emerge *Trichogramma* individuals (dark eggs) are introduced in central position. (b) The four types of experiments conducted, differing in how tubes were exposed to light and in their vertical orientation. The gray rectangles represent masks opaque to light. (c) From pictures taken every day, from D1 to D5, the parasitism rate of each patch in each tube is computed automatically using a color segmentation method.

In the experimental room, groups of 40 tubes were aligned, in a 2*20 grid. They faced a 40cm x 60cm light box, centered with respect to the grid. The light box was powered by a fluo lamp, delivering a constant 2,200 Lux of daylight illumination, homogeneous for all tubes, and with no measurable impact on tube temperature. The same 12:12 LD photoperiod was used as in rearing conditions. Four experimental treatments were conducted. First, in the “control” treatment, tubes were disposed horizontally, the light coming from above, all egg patches homogeneously lit (Fig. 1). This treatment did not provide any photactic or geotactic cues to discriminate the different egg patches, but was intended to capture a potential impact of other factors related to the experimental setup and ambient conditions. We conducted control treatments on a subset of eight populations spread over species (see Table 1). Second, for all 25 strains, we performed three treatments that introduced cues for oviposition preferences. In the “P” treatment, tubes were placed horizontally in the same manner, but each tube was masked on half of its length, so that half of egg patches were exposed to light, while the other patches were in darkness (Fig. 1b). Introduced parasitized eggs were thus exactly at the light/dark boundary and were after emergence offered a choice between the two sides. The masks were placed on the left halves of the tubes for 50% of them, and on the right halves of the tubes for the others, in alternating manner. This treatment allowed to investigate oviposition preferences with respect to light and phototaxis, but introduced no geotactic cue. In the “G” treatment, tubes were placed vertically, light still coming on them orthogonally, by effectively rotating the “Control” set-up by 90-degree clockwise (Fig. 1b). Tubes were not masked, so that all patches were homogeneously lit, but differed in elevation, introducing a geotactic cue to discriminate among them. Finally, in the “I” treatment, tubes were placed vertically as in the “G” treatment, but were masked from light on half of their length as in the “P” treatment (Fig. 1b). Masks were placed on the lower halves of the tubes, reproducing a natural set-up of “light” being associated with “upward”. This treatment introduces simultaneously photo- and geotactic components that discriminate patches, allowing to evaluate the interaction of the two factors.

For each population*treatment combination, 40 replicate tubes were used (i.e. one grid of 2*20 tubes described above). In one experimental session, i.e. on one particular week, all treatments (potentially including the control treatment) were conducted at once for one or two populations. This involved a total of 8 light boxes, 320 tubes, 3,200 egg patches, and about 2,500 adult *Trichogramma* individuals per week. All experiments were conducted in the same room by the same operator, in March-November 2018, randomizing the ordering of species over consecutive weeks.

### Image analyses

On the Monday following the beginning of an experiment (D1), parasitized eggs began to develop their typical dark coloration. As it takes five days for parasitized eggs to turn black, eggs turning black on Monday had been parasitized by the first adults that had emerged (on the previous Wednesday), eggs turning black on Tuesday had been parasitized the day after, etc. From Monday (D1) to Friday (D5), each block of 10 tubes was thus photographed every day in controlled lighting and exposure conditions using a hi-res digital camera, allowing to observe the dynamics of egg parasitism in each tube (see Fig. 1).

Each picture was processed through an ImageJ macro that automatically crops and numbers, from left to right, each of the 10 tubes. Each tube was then processed through another ImageJ/FIJI plugin, based on CODICOUNT^1^ (Schindelin et al. 2012; Perez et al. 2017), that identifies each of the 10 egg patches, and numbers them from left to right. For each patch, the plugin then computed the surface corresponding to parasitized eggs (dark in color) and to healthy eggs (pale in color), isolated from the background (cardboard). For each population and day, the plugin was trained by manually indicating samples of each class of pixels on a random subset (about 10%) of all pictures. The plugin then performs a discriminant analysis on color channels and determines an optimal segmentation, from which it returns a number of dark pixels (n1), corresponding to parasitized eggs, and a number of pale pixels, n2, corresponding to unparasitized eggs. The algorithm was then applied to all patches in all pictures, and for each we estimated the percent of eggs parasitized as n1/(n1+n2). The performance of the algorithm was tested by manually counting the number of healthy and parasitized eggs on a subset of pictures and comparing the results with the ones obtained with the automatic method. The parasitism rates obtained with the automatic method are very closely linearly related to those obtained from manual counts (r^2^ =0,95; see Figure S1 in Appendix; see also Perez et al. 2017; Dahirel et al. 2020; van Oudenhove et al. *in prep*.).

### Statistical analyses

Within a tube, egg patches were numbered from-5 to +5, negative indices corresponding to the left half side, and positive indices to the right side. Adult *Trichogramma* thus initially emerged between the -1 and +1 egg patches. In non-control tubes, by convention, positive indices were taken to correspond to the side for which an oviposition preference is expected, i.e. patches exposed to light in “P” treatments, patches in the upper part in “G” treatments, and patches in the light/upper part in “I” treatments.

Our protocol generated over 150,000 measurements of parasitism at patch level, from 3,000 tubes. Considering the large amount of data, and since our purpose was to characterize every population for its photo- and geo-tactic oviposition preferences and their additivity, we analyzed each population individually. To describe the pattern of parasitism in a tube on a given day, we computed two summary statistics: (i) the total rate of parasitism (all patches taken together, noted nbT), and (ii) the mean position of parasitism events (x), computed as the weighted-average of patch positions (from -5 to 5), each patch weighted by its rate of parasitism (fraction of eggs parasitized). A value of x close to zero indicates that eggs were parasitized indifferently on both sides of the tube, a negative or positive value on the contrary indicates a preference for parasitizing eggs on the corresponding side.

The value of x is an easily interpreted preference score: 0 indicates no preference, positive values indicate a preference in the expected direction, and negative values a preference opposite to expectations. We also considered the proportion of parasitized eggs on each side as an alternative preference score; results were similar, but x further takes into account how far on each side the individuals have been, and it is thus less influenced by the two centermost patches that are immediately adjacent to the release point. The scale of x is in numbers of patches: values -5 or +5 are the theoretical maxima corresponding to all parasitized eggs being in the farmost patches. A value of 2, for instance, means that parasitized eggs are on average located two patches away in the expected direction.

For each treatment, corresponding to 40 tubes of 10 patches, followed over five days for every population, we modeled the variation of x using a linear mixed model, using package lme4 in R version 4.x (R Core Team 2020). The 40 x values of the different tubes for a given treatment were weighted according to the square root of nbT. The square root is used because the precision of a mean estimate is inversely proportional to the square root of the number of observations (eggs n our case). Day (D1…D5) was used as a fixed quantitative effect (slope), Tube as a random effect, each observation weighted as explained above. Fixed effect estimates were used as summary statistics: the intercept (centered on Day 3) was used as a metric of the overall value of x, and thus of the strength of the preference. The slope over Day was used a metric as a metric of the change of x with time. For these two fixed effects, standard errors were obtained using parametric bootstrap with function Bootmer. From these, confidence intervals and pvalues (for non zero effect or difference) were obtained. To control for multiple tests, pvalues were adjusted using Holm’s method for each group of tests.

This procedure was first applied to control treatments in order to ascertain the absence of unforeseen experimental biases (8 populations tested), and then to the “P”, “G” and “I” treatments, to test for oviposition preference patterns (75 tests). For each population, we further compared the strengths of “P” and “G” preferences (25 tests), and tested for additivity of the two effects. To that end, from the “P” and “G” models, we predicted the intercept “I” treatments should have if effects were additive and the associated SE. We then tested for additivity by computing the difference of the predicted and observed “I” intercepts, its standard error, and the pvalue for it differing from zero (25 tests).

## Results

### Parasitism rates and control treatments

Our effective dataset (excluding controls) comprised 153,150 (15,315 tubes*day) estimates of parasitism rate at patch level. The overall parasitism rate varied across populations, ranging from 5% (pr002) to more than 20% (coret), with an average value of 11.8%, but no clear association with species was detected (see Figure S2). As expected, parasitism rate consistently increased over days, from an average of 9.2% on D1 to above 13% on D5. Averaged at tube level, parasitism rate followed a clear bimodal distribution, with a small mode close to 0 (presumably corresponding to failed or male-only hatching over the eight introduced parasitized eggs) and a larger mode, lognormal in shape, peaking at about 15% parasitism and reaching about 50% at most (Fig. S2). About 90% of tubes fell in the second mode, meaning they were actually parasitized. The 10% unparasitized tubes were omitted for subsequent analyses as they were uninformative with respect to oviposition preferences.

In the eight populations for which we conducted control treatments (see Table 1), the average position of parasitism (x) never significantly differed from zero, after correcting for multiple tests. Similarly, the mean position did not show significant systematic variations with Day, in any of the eight populations tested. This confirms that, intrinsically, the lighting setup and experimental tube configuration does not cause any systematic bias in the location of oviposition, as desired.

### Oviposition preference patterns

In the other treatments, however, most populations showed an oviposition preference, in one or the other direction (Fig. 2). Among the 25 populations, after correcting for multiple tests, 76% had a significant preference with respect to light (“P” treatments), 60% w.r.t. gravity (“G” treatments), Strain isa1072, from *T. cacoeciae*, stands out as the exception that had no preference at all. Note that this population had the lowest overall parasitism efficiency within *T. cacoeciae*, but not the lowest overall (Fig. S2).

**Figure 2:**
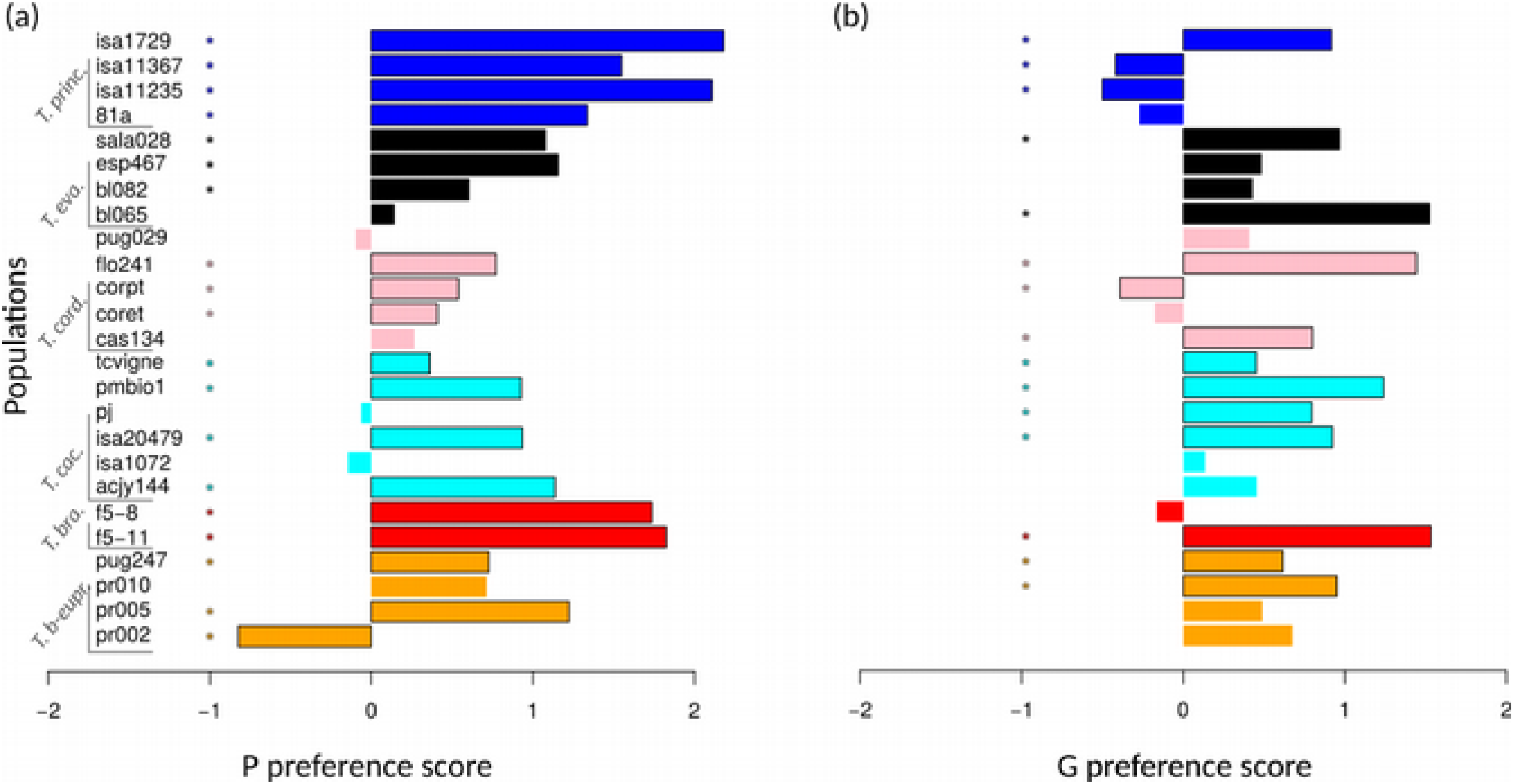
Preference scores for all 25 populations, in (a) “P” treatments (horizontal tubes with light/obscurity choice) and (b) “G” treatments (vertical tubes with light everywhere). Positive scores indicate preference in the expected direction (for light and upward positions, respectively). Values significantly different from zero after correction for multiple tests are highlighted with an asterisk and a dark frame. Colors correspond to different species (see Table 1).

The vast majority of populations had the expected preference for ovipositing in well-lit areas rather than in darkness, only one in 19 (pr002) having the opposite preference (Fig. 2a). In contrast, three populations (on the 15 that had a significant vertical preference) had a preference for ovipositing in lower parts of the tube, contradicting the expectation (Fig. 2). These are corpt (species *T. cordubensis*), isa11235 and isa11367 (both from species *T. principium*). Example distributions of parasitism for population isa11235 are shown in Fig. S3.

Preference for light therefore seems a much more consistent characteristic in *Trichogramma* than preference for going up. This is confirmed by the fact that, regardless of direction, the absolute score of vertical preference was weaker than the score of preference for light in a majority (68%) of populations. Restricting to significant differences between the two scores (10 populations, after correction for multiple tests), the picture is similar, with 70% of differences being in the direction of a stronger light preference.

Finally, 44% of populations had a significant preference in both “P” and “G” treatments. This corresponds to the frequency expected if the two traits were distributed independently across populations. Consistent with that, there was no detectable correlation between the strength of the two preferences across populations (Spearman rank correlation; rho=-0.07; p=0.74).

The preference scores discussed above were not static in time, but systematically varied with day: they tended to be strongest in the first days and to revert to zero as days passed (Fig. 3a). Across all populations and treatments, after correcting for multiple (75) tests, the mean position of parasitism x was significantly affected by Day in 68% of cases. All significant slopes were negative, with no exception, and overall, there was a strong negative correlation between the strength of the oviposition preference x and its slope w.r.t. Day (Fig. 3b; Spearman rank correlation; rho=-0.58; p=7.6e-8). This indicates that the stronger a preference was, the stronger was its tendency to decline over consecutive days. All this is strongly consistent with a resource depletion effect: as individuals preferred to lay eggs on one side, they depleted the amount of available host eggs on that side. This gradually encouraged them to overcome their initial preference and go explore the other side where host eggs are still plentiful. Such a pattern was not necessarily expected, considering the very large number of host eggs present in the tube, and consequently the parasitism rates that remained fairly low overall (see Fig. S2). In specific patches though, parasitism rates could approach 50% after two or three days (Fig. S3).

**Figure 3:**
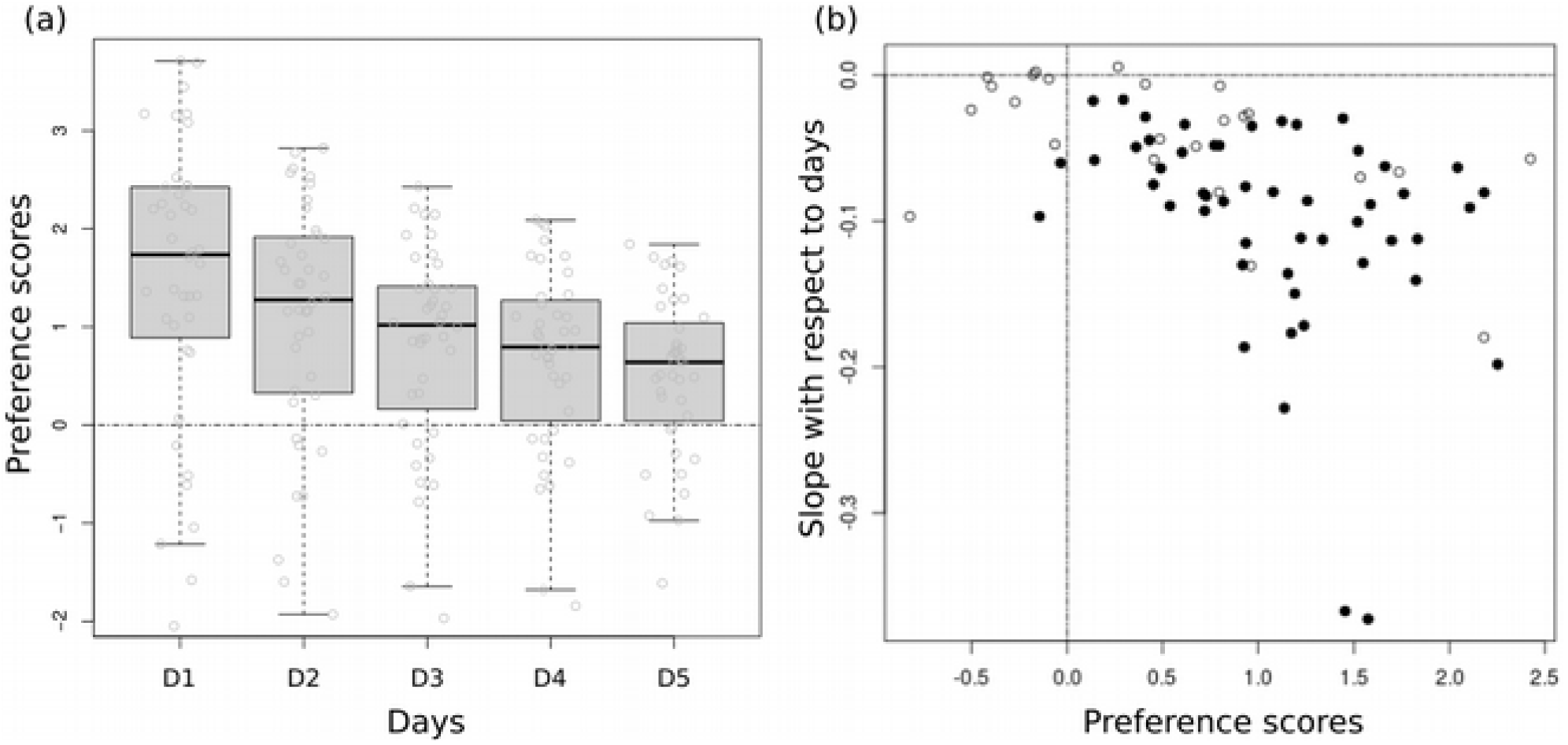
Temporal dynamics of oviposition preference. (a) Distribution of x values (one dot per tube; 40 tubes) as a function of Day, in the example of population acjy144, “P” treatment. (b) Correlation, over all populations and treatments (75 points), between the preference score (x axis) and its change over days (y-axis). Black dots indicate population/treatment combinations for which the slope was significantly different from zero, after correcting for multiple tests.

### Additivity of orientation cues

96% of populations (all but isa1072) showed a significant preference when both gravity and light were offered as cues (“I” treatments; Fig. 4a). Preferences were thus, as expected, stronger when two cues were simultaneously present. But were the effects of the two cues simply additive, or did they interact in a more complex way? After correcting for multiple tests, only 8% of populations (two out of 25) had a significantly non-additive response to the two oviposition cues: corpt (*T. cordubensis*) and isa1729 (*T. principium*).

**Figure 4:**
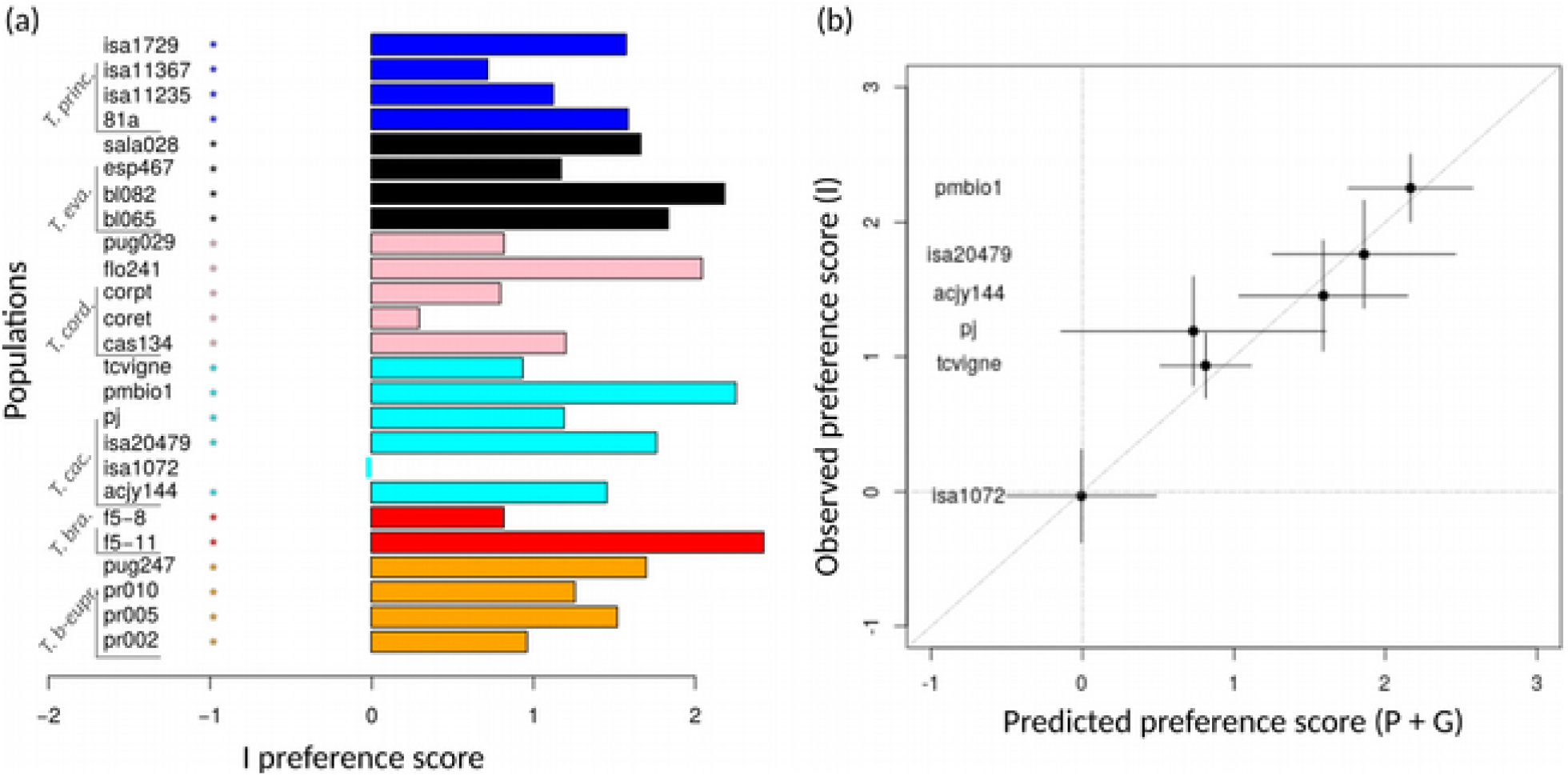
Interactions of photo- and geo-tactic oviposition preferences. (a) “I” preference scores for all 25 populations. Values significantly different from zero after correction for multiple tests are marked with a black box. (b) Correlation between predicted (x axis) and observed (y axis) preference scores for the six populations of species *T. cacoeciae*. Horizontal and vertical bars are 95% confidence intervals.

Population corpt was remarkable in having a preference for ovipositing in lower positions in “G” treatments, and a weak preference for light in the “P” treatment (Fig. 2). It had, however, a significant preference to lay in the light and upper egg patches in the “I” treatment, causing the non-additivity pattern (Fig. 4). This suggests a synergistic action in this population: preferences w.r.t. to light or to gravity were individually relatively weak or atypical, but when signals were combined in the usual way (light being up), they elicited a clear preference in the expected direction.

Population isa1729 had a simpler sub-additive preference pattern: it had strong and significant preferences for laying eggs in well-lit and uppermost egg-patches, but in the “I” treatments, the resulting preference pattern was weaker than expected, even though in the expected direction. This is suggestive of a saturation effect: both cues were individually sufficient to elicit a strong preference, and the latter could not be increased much further in the presence of the two. This might be related to the depletion effect highlighted above: the stronger the preference, the faster resource depletion is, and thus the greater the repulsive effect that tends to counteract the initial preference (see also Figure S3).

Apart from these two particular cases, the overall pattern is therefore one of independent actions: the contributions of the two cues appear as remarkably additive. This is best illustrated within species *T. cacoeciae*, of which we had five contrasted populations. The preference scores observed in treatments “I” are almost exactly as predicted by summing the two scores observed in the “P” and “G” treatments (Fig. 4b).

### Connection to vegetation strata and species

Considering the general consistence of “I” treatments with the other two, we included all three treatments together (i.e. 120 tubes per population) and performed a PCA to have a synthetic representation of all populations in the plane (Fig. 5). As expected, the first two components explained most (94%) of the variance. The first axis tends to quantify variation in overall preference strength, discriminating populations with strong preferences from those with weak or no preferences. The second axis rather discriminates populations along the relative strengths of photo-versus geo-tactic preferences.

**Figure 5:**
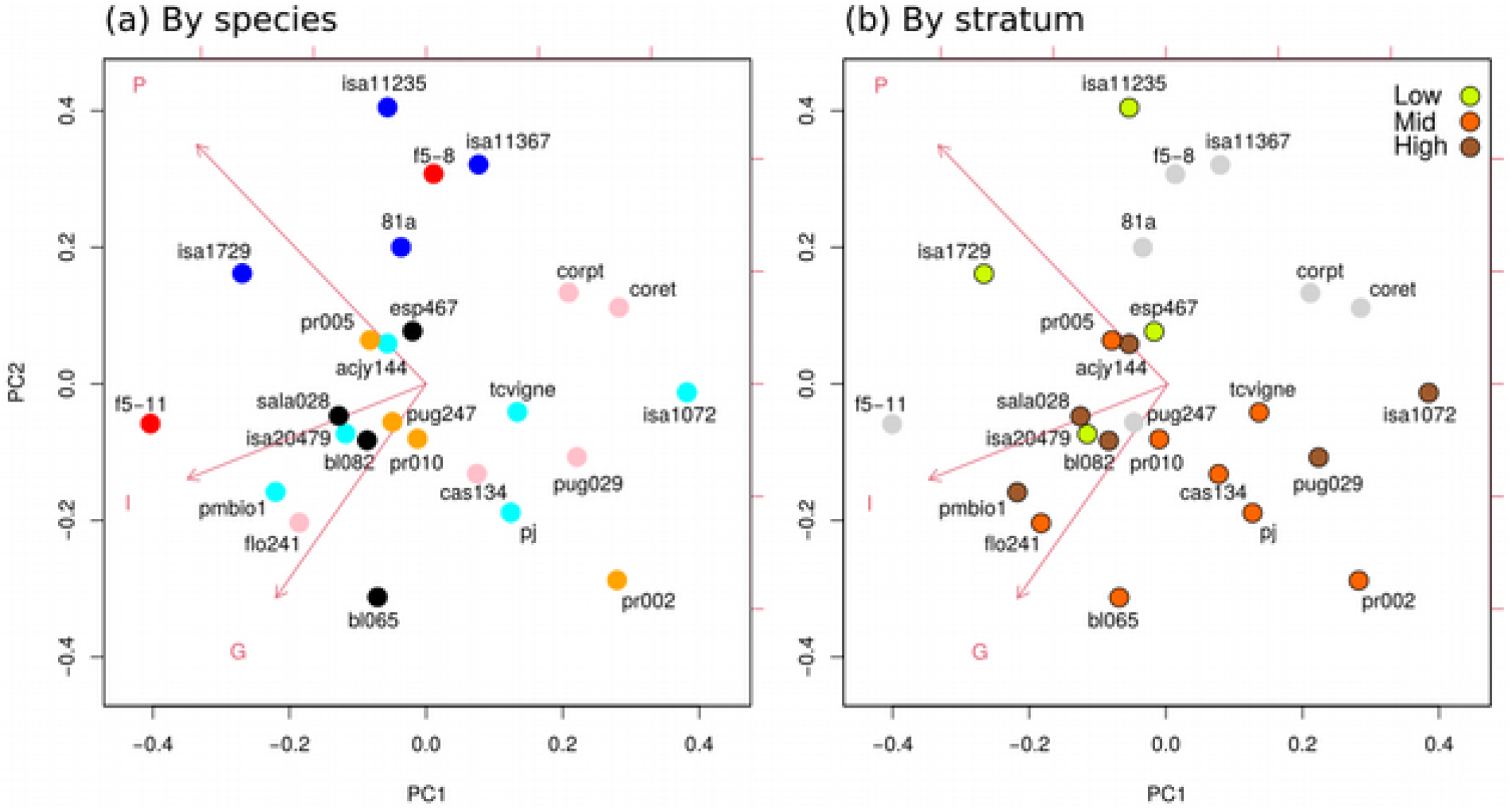
PCA representing all 25 populations based on their three oviposition preference scores (“P”, “G” and “I” treatments). The three variables are represented by the three red vectors. PC1 and PC2 captured 51% and 43% of the total variance, respectively (see Table S1 in S.I.). Populations are colored according to (a) species (see Table 1 and previous figures for color code) and (b) the vegetation strata in which they were sampled. In (b) grey circles are populations without stratum data available.

Grouping populations by species (Fig. 5a), it is obvious that there is an important level of intraspecific variation. Species show considerable intraspecific variation along axis 1, populations thus differing in their level of oviposition preference. In general, variation is smaller along axis 2. Species thus tend, qualitatively, to be consistent in their preference patterns. This is most visible in *T. cacoeciae*, that differs importantly along axis 1, but very little along axis 2: all populations share a common pattern of preference for light and preference for going up (see Fig. 2). Even though we must be cautious in our interpretations considering the relatively low numbers of populations per species, at species level, the most striking pattern is that *T. principium* and *T. brassiceae* populations form a distinct cluster characterized by a very strong preference for light. This cluster can be delineated in Figure 5a by drawing a line perpendicular to the “P” vector. Consistently, the “P” preference score significantly differs across species (Kruskal-Wallis test; p=0.016). *T. principium* is further characterized by a tendency to have weak, and often slightly negative, geotactic preferences, unlike most other species.

Regarding vegetation strata, interpretations are even more tentative because of the partial information available. Still, the prediction that populations sampled in lower strata should have greater propensity to oviposition in lower positions and greater preference for light received some qualitative support (Fig. 5b). Indeed, the four populations sampled in the lower stratum have on average stronger preferences for light and never have very strong preferences for ovipositing in uppermost patches. In contrast, the two upper strata showed no detectable differentiation. Consistently, restricting the analysis to populations with known vegetation stratum, “P” preferences significantly differed across vegetation strata (Kruskal-Wallis test; p=0.035), and so did the score of PCA component 2 (p=0.028). This pattern is clearly not independent of the one discussed above, as two of these populations belong to species *T. principium*. Taken together, these results suggest that species may be distributed differently across vegetation strata, possibly with a corresponding variation in their oviposition preference traits.

## Discussion

As expected, most *Trichogramma* populations and species were found to have a preference for light versus shade, and for upper positions versus lower positions (Fig. 2). However, our study revealed ample quantitative, and sometimes qualitative, differences across populations and species. The overall picture was indeed quite clear when both orientation cues are offered simultaneously, as is done most often, and as probably typically occurs in natural conditions: virtually all populations then harbored the expected preference, with only one exception (Fig. 3). This was very much less the case when the two cues were experimentally dissociated. Not all populations preferred light, with one counterexample preferring shade. The picture was even dimmer for geotactic preferences: only 60% of populations harbored a significant preference in the absence of light as a consistent orientation cue (Fig. 2b). This indicates that, overall, photo-related preferences were stronger overall than gravity-related preferences. It is of course difficult to compare the strength of the stimulus in the two types of experiments, because light and gravity are so different, and because we lack detailed knowledge on the neuro-sensory systems involved in these insects. However, the levels of light we used were not extreme (corresponding to a typical overcast day), and the presence of a 12:12 photoperiod implied that the light stimulus was only available half of the time, whereas gravity was always available. These results therefore suggest that in these insects, gravity is overall less important a factor for microhabitat selection and oviposition preference than light is.

The two traits were otherwise independently distributed, and were almost perfectly additive in their contributions to the oviposition preference patterns. There is therefore no indication of an underlying genetic variation in overall “choosiness” that would simultaneously impact both photo- and geo-tactic oviposition preferences. Similarly, there appears to be no negative trade-off like association or other interaction between the two components. A consistent association of photo- and geo-taxies is not necessarily expected in natural conditions, since the distribution of light and shade can easily occur independently of vertical position. Similarly, the mechanisms of photo- and gravi-perception, even though imperfectly known in insects, are probably distinct and could well be largely independent. Overall, this suggests that the two types of preferences are not tightly associated and could evolve or be selected for independently if needed, at least to some extent. The only exceptions to this were two populations, corpt and isa1729, that responded in the presence of both orientation cues in a way that could not be predicted from the two single-cue experiments. These two cases differed in the type of non-independence however. In the first case (corpt), the pattern could be indicative of a synergistic effect, whereby light preference is not fully expressed in the absence of any gravity-related cues. In the second case (isa1729) on the contrary, the single-cue oviposition preference patterns were already so strong that when combined (“I” treatments) that they failed to reach the level predicted when combining the two.

The latter result is well in line with another general conclusion from our experiments: the oviposition patterns we observed were innate but very plastic. In particular, they were very dynamic in time, with a general tendency to have strong preferences initially (on the first day), followed by a gradual decline over the following days, to the point that the final pattern of parasitism (on Day 5) could show no significant preference overall. Among the various possible interpretations, the one we consider most likely is that this is caused by host depletion: a strong initial preference causes parasitism to be concentrated in the preferred parts. It is well-known that individual *Trichogramma* females can detect host eggs that have already been parasitized and can adjust behavior accordingly (Wajnberg et al. 2000). Therefore, females would soon be deterred away from preferred areas for lack of unattacked eggs, causing a reduction, or even an inversion, of the effective oviposition patterns. This result was not necessarily expected in our experimental conditions, considering the supply of host eggs was large relative to the number of females and, consequently, that parasitism rates remained fairly low overall (Figure S2). However, at the level of individual patches, parasitism rates could get close to 50% after two or three days (see Figure S3), so that depletion of preferred patches might suffice to cause the observed trend. Alternatively, the latter could be explained by habituation or aging effects, whereby initial preferences are gradually relaxed. Indirect interactions through host parasitism, conspecific chemical cues or direct inter-individual interactions may also contribute to the dynamics of oviposition patterns. In any case, these results show how plastic, and in particular how sensitive to host distribution and to intraspecific competition, oviposition patterns are, even though they are expected to be almost universal and quite strong in the short term. For this reason, realized oviposition patterns may be challenging to extrapolate from short-term individual behavioral observations, and they might be easily obscured by plasticity if studied with insufficient resolution. This underlines the need to study oviposition preferences over the course of time, taking into account the dynamics of parasitism.

Host searching in our experimental tubes most likely resulted in random search, coupled with general orientation cues (related to light and gravity), plus short-range visual cues. Indeed, the *Ephestia kuhniella* eggs we used emit little odorant molecules and were devoid of moth scales or other Lepidopteran kairomones. Considering that, even in field conditions, evidence for odor orientation in *Trichogramma* is mixed (Gardner & Hoffmann 2020), it is unlikely to have played an important role in our setting. Note that patches of host eggs were separated by a distance (about one centimeter) larger than the typically reported reactive distance of *Trichogramma* (less than four millimeters in this type of conditions; Bruins et al. 1994). Therefore, the movement of *Trichogramma* in between egg patches, besides random search and thigmotaxis (i.e. tendency to follow physical boundaries of the environment), must have relied largely on other orientation cues, in particular light and gravity as we offered.

The effects of light and gravity we reported on oviposition patterns probably integrated a number of different elementary behavioral processes over several days. These can range from phototaxis and geotaxis *stricto sensu* at short temporal scales, to associative learning effects on longer timescales, going through various types of ortho- and klino-kinesis (changing nature of movements in response to stimulus). It would be interesting to dissect the relative contributions of such behavioral processes, but this is beyond the scope of this work and would require more detailed behavioral assays. Here our interest is in the resulting parasitism distributions, and we use “photo- and “geotactic” in a less formal sense to describe the observed patterns of oviposition preference in relation to light and gravity.

The taxonomy of *Trichogramma* is notoriously complicated and is undergoing rapid changes, as new tools from integrative taxonomy are being employed (Consoli et al. 2010). Species boundaries are often difficult to delineate, and even when clearly delineated, it is likely that an important geographical or host-related diversity remains concealed, waiting to be discovered. This is not an easy task, for the field biology of *Trichogramma* is very poorly known, and sampling of natural populations is labor intensive. In these conditions, many studies deal with only one or a few laboratory strains per species, and we still have very little idea of the population structure and intraspecific variability in nature. Morphological differences between species are minimal, and most species are virtually indistinguishable. As a consequence, inter-specific differences cannot be easily related to variations in other traits, such as body size, flight capacities or host ranges, as one might envision. This study is an effort in the direction of better characterizing *Trichogramma* diversity, since we sought to characterized several recently sampled and well-sourced populations per species. The first obvious conclusion is that intraspecific variability is very important. This makes sense from an ecological perspective: behavioral traits related to microhabitat preference are expected to be very evolutionarily labile, as they are key fitness components for adaptation to the local habitat conditions and to particular host species and organs. In particular, it is known that Lepidopteran females have quite specific preferences at oviposition, with respect to to plant parts, height or the upper versus lower sides of leaves (e.g. Hussein & Kameldeer 1988; Conti & Bin 2000; Spangler & Calvin 2001; Singer 2015).

An important related question is the impact of laboratory rearing conditions on the phenotypes we have studied. Even though most populations were sampled only a few years before experiments, our populations might have undergone significant evolutionary changes in the lab, especially considering their short generation times (less than one month). To what extent is the variability observed here representative of the natural variation? Did rearing conditions bias the oviposition preferences in certain ways? These questions are hard to answer, but given our rearing protocols, one could hypothesize that laboratory populations would lose many of their oviposition preferences over time, for they have no adaptive value in the lab. The only element we can put forward is a comparison between our “old” laboratory populations, that have over 20 years, and the “young” ones, that have only a couple of years of lab adaptation (see Table 1). As it appears, no systematic difference emerged. The example of *T. cacoeciae* is the most telling: two out of six populations (pj and tcvigne) were “old”. Yet, as is most visible in Figure 3b, they were in no way atypical. This would suggest laboratory evolution did not play an important, or at least predictable, role, but of course dedicated studies will have to be conducted to address this question.

At species level, the most salient pattern was a tendency for two species (*T. principium* and *T. brassicae*; 6 populations in total) to be characterized by very strong positive phototactic preference (Fig. 5a). In terms of habitat, the populations we had for *T. principium* were more associated with the lower strata (herbaceous layer), and populations from this strata were consistently never among those with lowest phototactic preferences (Fig. 5b). There might thus be a tendency for populations/species from the herbaceous layer (such as *T. principium* for instance) to be associated with stronger preference for ovipositing in light-exposed areas rather than shade. Such an ecological differentiation could make sense from an adaptive perspective, since low herbaceous plants and grasses are often exposed to direct sunlight, whereas shrubs and trees cast more substantial shade within their crowns. Unfortunately, our sample size (25 populations), though decent, is still insufficient to draw any firm conclusion, especially considering the partial ecological data in our possession. At best, we can draw some tentative hypotheses and encourage further investigations along those lines.

The phenotyping protocol introduced here offers a way to characterize a significant number of strains or populations with high replication power, moderate cost, and minimal workload. Indeed, the use of automated imaging techniques to individualize pictures of individual tubes and host-patches and evaluate their level of parasitism greatly diminished the time required for these tasks, allowing for greater throughput. It was estimated that counting the number of parasitized and healthy host eggs using the usual manual procedure is as much as 12 times longer than the method used here. Furthermore, the automatic method also makes the counting process more objective and more reproducible. The macro codes used for counting are exported and can be subsequently reused at any time to replicate exactly the same counting protocol, irrespective of the operator’s identity or condition (Powers & Hampton 2019). The variability of the counting processes can furthermore be quantified and formally taken into account in statistical analyses if needed (see e.g. Dahirel et al. 2020). This approach has already been used in several projects with satisfying performance, and there is still ample room for improvement. We believe it has great potential to facilitate and increase the scope and power of a variety of experimental investigations in *Trichogramma* and similar biocontrol agents. Going back to the question at hand in the present article, these methods provide a platform to achieve even greater sample sizes and reach the necessary power to test for the trait-habitat associations suggested above in a more decisive way.

The intra- and inter-specific variability reported here in *Trichogramma* has important practical consequences for biological control. Agricultural pests are known to differ in their microhabitat preferences and/or in the parts of the growing plants on which they oviposit most (e.g. Conti & Bin 2000; Gardner & Hoffmann 2020). As a consequence, strains of *Trichogramma* that have oviposition preferences matching those of the target pests should be preferred, and could provide better control performance. In the context of inundative releases of *Trichogramma*, our results suggest that strains may be selected, beyond usual performance characteristics, for oviposition preferences that match the favoured oviposition sites of the target pest, and attributes of the target crop, such as plant height. Beyond that, mixtures of strains with different microhabitat selection preferences might provide a better overall coverage of the host plants, together with reduced intraspecific competition in the preferred parts. Such diverse mixes might thus achieve better overall performance, compared to any particular monomorphic inoculum, as we can expect in trophic systems (Schmitz 2007; Bruno & Cardinale 2008; Kering et al. 2019). We suggest photo- and geo-tactic preferences as studied here provide operational phenotypic traits, and traits quite distinct from those more usually studied, for research and selection purposes.

## Acknowledgments

This research was funded by the French ANR (project TriPTIC) and by INRAE (project SPE-PHENOMENE). Thanks to the EP-Coll Biological Resource Center for biological material. The collection/BRC EP-Coll is a part of BRC4Env, the pillar “Environmental Resources” of the Research Infrastructure AgroBRC-RARe. Version 4 of this preprint has been peer-reviewed and recommended by Peer Community In Zoology (https://doi.org/10.24072/pci.zool.100008).

## Conflict of interest disclosure

The authors of this preprint declare that they have no financial conflict of interest with the content of this article. Vincent Calcagno is one of the PCI Zool recommenders.

## Data availability statement

The entire raw dataset and associated scripts and informations are available online from https://doi.org/10.5281/zenodo.4665806.

## Supplementary Information

**Figure S1:**
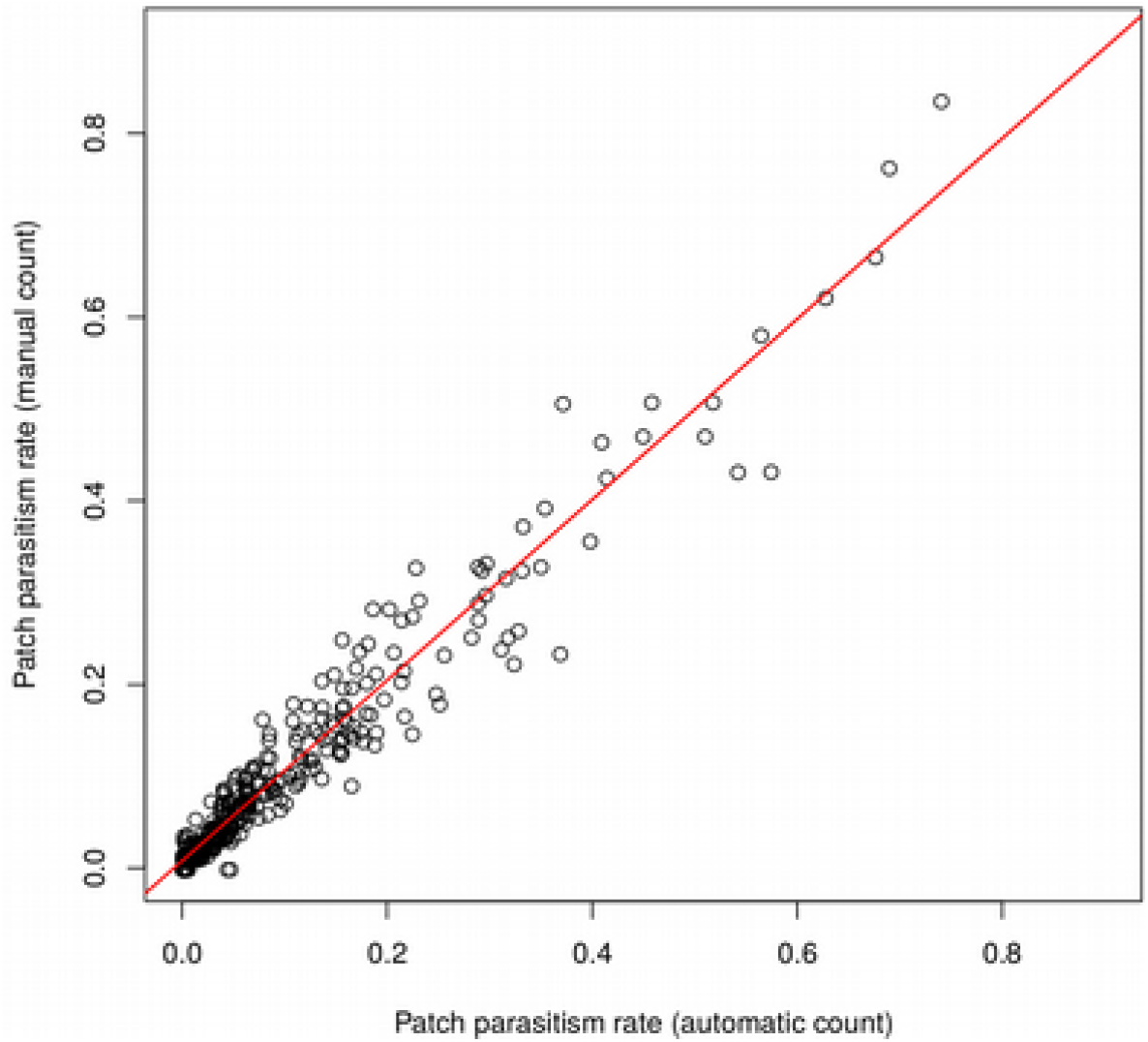
Correlation between automatic and manual estimates of patch parasitism rates. A subset of 400 experimental patches (200 from population acjy144, and 200 from pr002, both in “H” treatments) were manually counted. The parasitism rates obtained from the automatic counts (x-axis) and those from the automatic count (y-axis) are very closely related. The slope of the linear regression model shown in red is 0.983, its intercept is 0.008, and the coefficient of determination r^2^ is 95%. For comparison, 40 patches (20 from each population) were also manually counted by two different operators: the linear regression between the two sets of manual counts had slope 1.03, intercept -0.07, and r^2^=99%.

**Figure S2:**
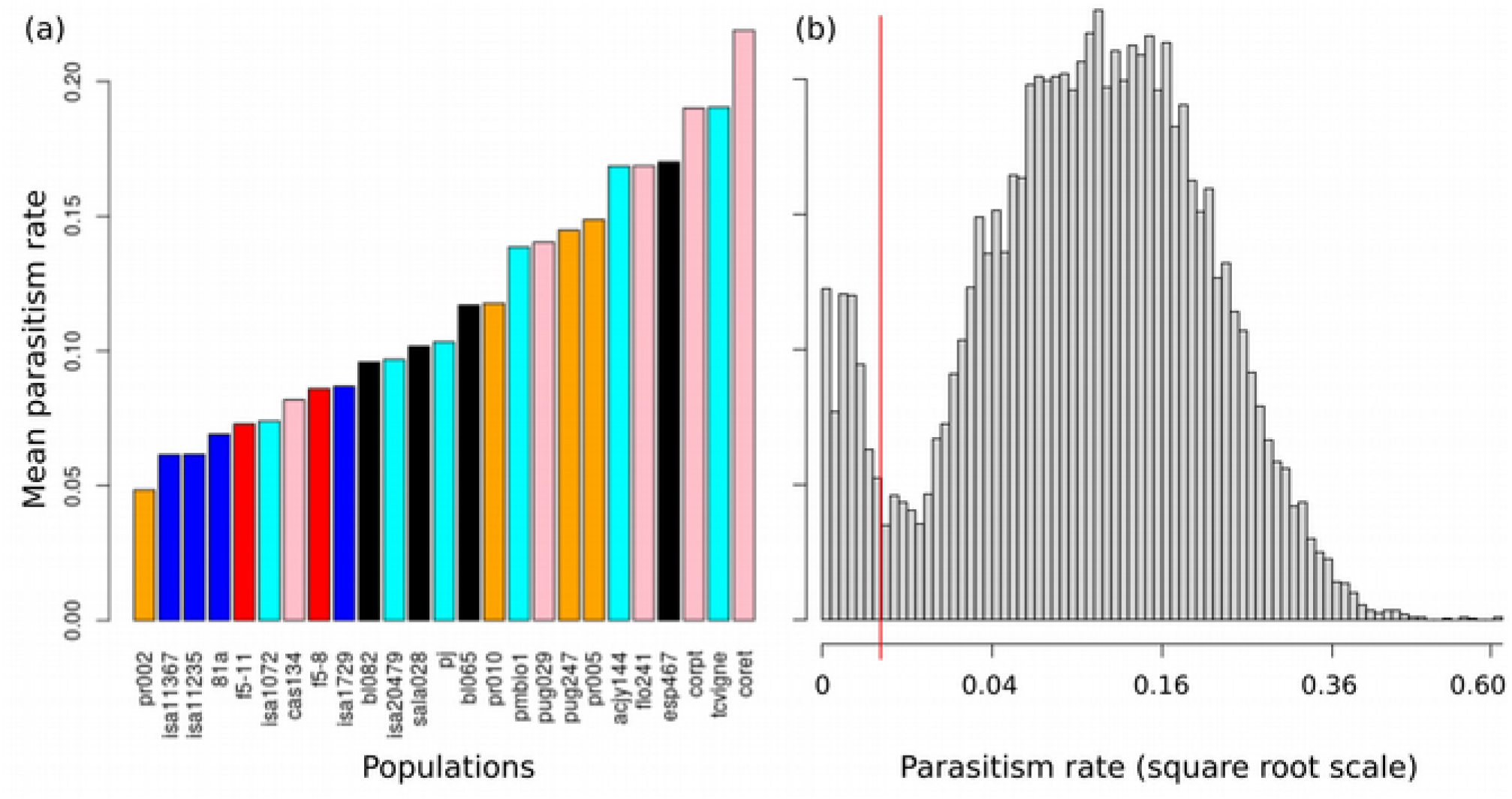
(a) Mean overall parasitism rate, over all patches, treatments and days, for each of the 25 populations studied. Populations are sorted from lowest to highest parasitism efficiency, and are color coded according to species membership (see Table 1). (b) Distribution of parasitism rate per tube, across all populations, treatments (omitting controls) and days. The leftmost peak (on the left of the red vertical line) was omitted from the analysis of oviposition preferences for lack or absence of parasitism. The right peak is almost normally distributed on the square root transformed scale. Parasitism rates at patch level are presented in Figure S3,

**Figure S3:**
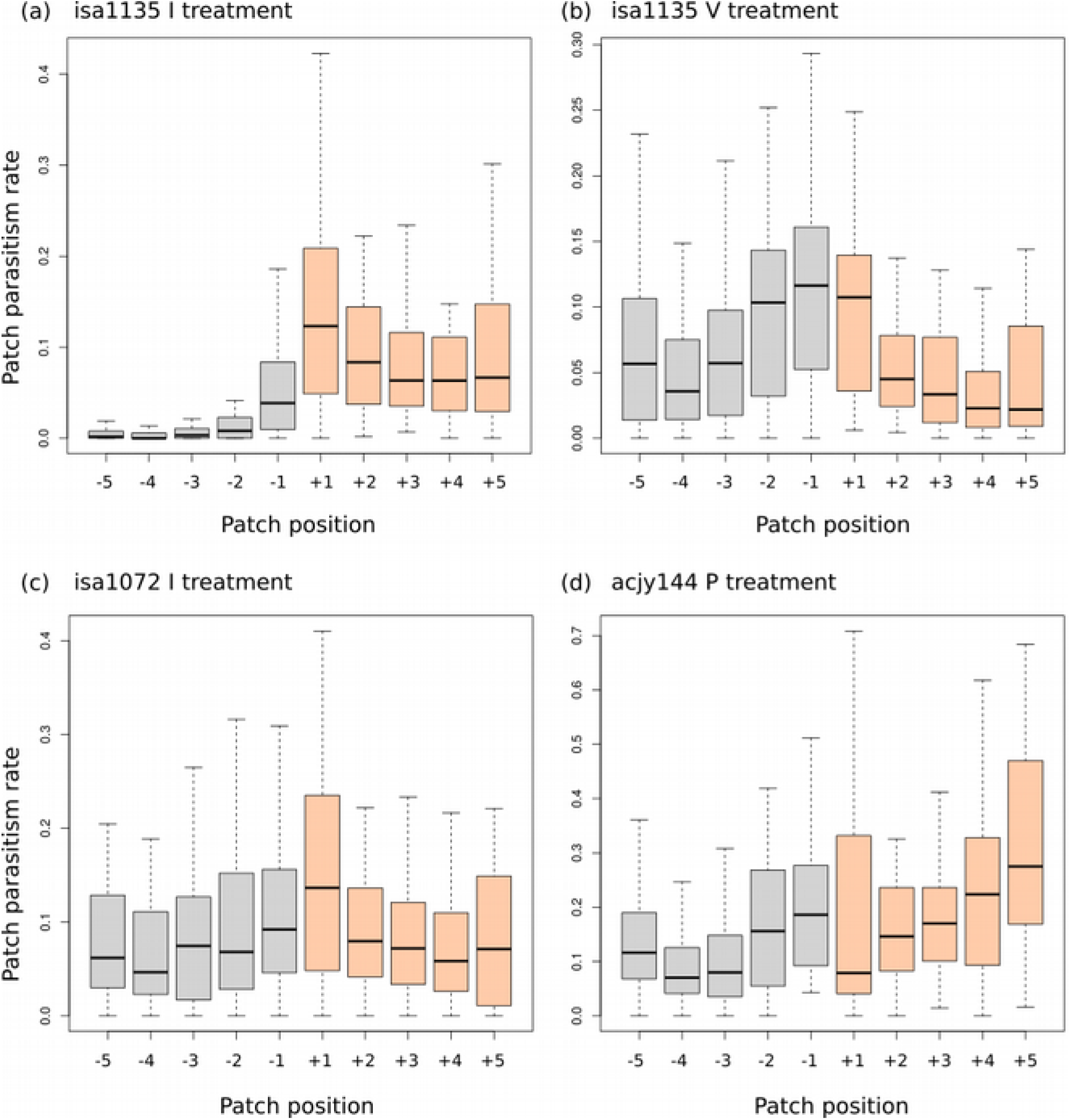
Examples of the distribution of parasitism rate at patch level, along the experimental tubes. Top panels: population isa1135, on Day 2, in the “I” treatment (a) and the “V” treatment (b). Remark that this population features a clear, atypical, preference for ovipositing in the lower parts. Panel (c): population isa1072, on Day 3, in the “I” treatment. Note the atypical absence of any particular preference. Panel (d): population acjy144, on Day 3, in the “P” treatment. This is the example shown in Figure 3a, that features a very pronounced “relaxation effect” over consecutive days. Positive patch numbers (in orange) are on the side for which a preference is expected (well-lit and in upward positions in “I”, upward positions in “G”, well-lit positions in “P”).

**Table S1:** Loadings for the principal component analysis presented in Figure 5. The PCA was performed using function prcomp in R, based on estimates from the linear mixed-models.

|  | PC1 | PC2 | PC3 |
| --- | --- | --- | --- |
| Proportion of variance | 0.5111 | 0.433 | 0.05554 |
| “P” loading | -0.6310106 | 0.7143917 | -0.3024402 |
| “G” loading | -0.4131820 | -0.6394478 | -0.6483727 |
| “I” loading | -0.6565868 | -0.2841671 | 0.6986722 |

## Footnotes

1 https://www6.paca.inrae.fr/institut-sophia-agrobiotech_eng/CODICOUNT

## Notes

https://doi.org/10.5281/zenodo.4665806

### Competing Interest Statement

The authors have declared no competing interest.

### Summary of Updates

Version 4 of this preprint has been peer-reviewed and recommended by Peer Community In Zoology (https://doi.org/10.24072/pci.zool.100008)

